# SRSF shape analysis for sequencing data reveal new differentiating patterns

**DOI:** 10.1101/161448

**Authors:** Sergiusz Wesolowski, Daniel Vera, Wei Wu

## Abstract

**Motivation:** Sequencing-based methods to examine fundamental features of the genome, such as gene expression and chromatin structure, rely on inferences from the abundance and distribution of reads derived from Illumina sequencing. Drawing sound inferences from such experiments relies on appropriate mathematical methods to model the distribution of reads along the genome, which has been challenging due to the scale and nature of these data.

**Results:** We propose a new framework (SRSFseq) based on Square Root Slope Functions shape analysis to analyse Illumina sequencing data. In the new approach the basic unit of information is the density of mapped reads over region of interest located on the known reference genome. The densities are interpreted as shapes and a new shape analysis model is proposed. An equivalent of a Fisher test is used to quantify the significance of shape differences in read distribution patterns between groups of density functions in different experimental conditions. We evaluated the performance of this new framework to analyze RNA-seq data at the exon level, which enabled the detection of variation in read distributions and abundances between experimental conditions not detected by other methods. Thus, the method is a suitable supplement to the state of the are count based techniques. The variety of density representations and flexibility of mathematical design allow the model to be easily adapted to other data types or problems in which the distribution of reads is to be tested. The functional interpretation and SRSF phase-amplitude separation technique gives an efficient noise reduction procedure improving the sensitivity and specificity of the method.

## 1. Introduction

Second-generation sequencing technologies, such as Illumina sequencing, has allowed researchers to discover fundamental features of genomes and their regulation, organization, and dynamics. For example, sequencing experiments that examine the dynamics of transcription of genomic DNA into RNA (RNA-seq), involve the isolation of RNA from populations of cells which are experimentally processed and sequenced. The generated sequences are often mapped to a reference genome to identify the genes from which the sequences originated. The quantification of sequences mapped to each gene are then used to estimate the level to which each gene is expressed. These gene expression estimates are often compared between experimental conditions to make inferences about gene expression differences. This sequence-map-quantify-compare paradigm is the basis for many functional genomics experiments, including ChIP-seq, DNase-seq, MNase-seq, and Bisulfite sequencing. The data provided by these experiments are in the form of genomic coordinates to which the reads are predicted to be derived from, which number on the order of tens of millions of observations. Because of various biological and technical aspects of these experiments, these read distributions have proved difficult to model (Hayer et al., 2015). While there have been numerous attempts to accurately model these data, nearly all involve reducing data in a form that summarizes read counts over defined genomic regions, which discards or significantly reduces information on the spacial distribution of reads.

In the case of RNA-seq or ChIP-seq, which is typically focused on examining discrete genomic units of genes, these read distributions are generally reduced by summarizing the number of reads that map to each gene. This approach discards information about the spacial distribution these reads derive relative to the gene or exon (Figure 1). While this simplifies the complex data and is more easily modeled, it also loses information that may provide insight on gene expression dynamics that are associated with the shape of the distribution of reads, (e.g as alternative splicing and variation in transcription start and termination sites, differential exon usage). This method also suffers from inaccuracies introduced by the presence of overlapping genes, which may cause inaccurate counting of one gene caused by the expression of another overlapping gene. Several statistical models have been developed to attempt to address these issues, including DESeq2, DEXSeq, Cufflinks, Limma, BaySeq, EBSeq, (Love et al., 2014; Anders et al., 2012; Li et al., 2015; Trapnell et al., 2012, 2013; Law et al., 2014; Hardcastle and Kelly, 2010; Leng et al., 2013).

**Figure 1:**
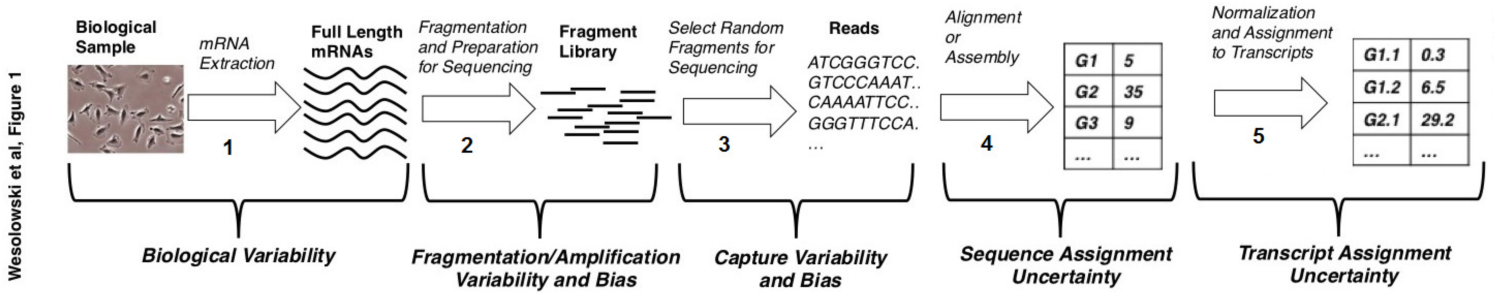
Next Generation Sequecning (NGS) workflow schematic with highlighted possible variability biases. The focus of the methods in this article is on redefining steps 4 and 5.

Here, we present the SRSFseq, a new framework for analyzing genomics data based on second-generation sequencing. The framework interprets the distribution of read alignments across the genome as shapes. This approach takes into account information provided by the base-level distribution of the mapped reads in order to examine variability in the shapes of the read densities over genomic regions. It takes into account the relative read abundances and the differences in read density profiles. We show how this framework can be used to identify new differential expression behaviors and successfully supplement the results established by state-of-the-art, count based methods.

In the following sections we show the functionality of the SRSFseq on an example of exon level differential expression analysis. The functional interpretation of the model allows us to use the phase-amplitude separation (Srivastava et al., 2011) which accounts for additional levels of the noise and the data normalization. We utilize the functional F-test (Zhang, 2013) to determine the differential expression and compare the results with selected popular methods: Cuffdiff, DESeq2 and Limma-voom. Next we propose an alternative extension of the SRSFseq in application to the shift and the shape change detection in MNase-seq data for nucleosome positioning.

## 2. Methods

In order to model the distribution of read alignments, first we obtain the read densities of a specific genomic region of interest (*G*). This region is assumed to be common among different samples, e.g. exon, gene, transcription starting site. The read densities, from now on, are treated as the basic unit of information for further modelling. In this work to obtain read densities, we are using standard kernel density estimator applied to coordinates of the mapped reads as seen in an example in Figure 2. Throughout this paper we will refer to this step as the filtering. The choice of the density estimation technique, may bear significant influence on what features of the Next Generation Sequencing (NGS) data are to be extracted. As a consequence of the filtering step, the data is automatically normalized.

**Figure 2:**
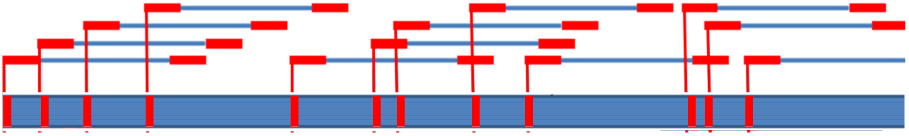
Obtaining the point pattern data from the mapped reads. The leftmost coordinate of each read on the reference genome is reported.

Summarizing, our approach moves the area of modeling from vector-valued variables, used in most of the available methods, to infinitely dimensional space of read densities. The mathematical complexity level is higher, but it is necessary to benefit from the advanced shape analysis modeling tools to unlock the full potential of the Next Generation Sequencing.

### 2.1. Functional ANOVA for read densities

The density normalization (filtering) is essential, as we want to focus to uncover new information stored in the NGS results. The density normalization allows us to mod-out all differences coming from discrepancies between number of mapped reads as well as make the data comparable between experimental samples. As our normalized data no longer depends on read counts, we expect to detect different information encoded in the NGS than the count-based methods. We confirm this hypothesis in the section 3.

In general SRSFseq is suitable to compare and model any point patterns arising from mapping NGS reads to a reference genome. For sake of clarity we focus on exon level differential gene expression and RNAseq experiments, but we want to emphasize that the methods described below can be applied to any NGS output as long as it consists of mapped reads over known genome.

In the example we aim to be able to compare the gene expression patterns between *j* = 1, *… k* conditions over a genomic region of interest (in our case, exon). In our approach a gene is differentially expressed if at least one of it’s exons is statistically significantly differentially expressed and different gene isoforms are treated as different genes.

To quantify the difference between conditions we utilize the Functional ANOVA F-Snedecor test with null hypothesis:

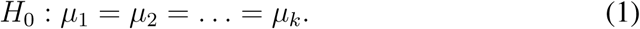

The test statistic is a ratio of the sum of squared distances between and within conditions scaled by their degrees of freedom.

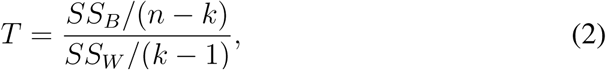

where *k* represents the number of conditions tested and the *SS*_*B*_ and the *SS*_*W*_ are the sums of squared distances within and between conditions, as defined in the ANOVA statistics, but utilizing the *L*^2^ norm:

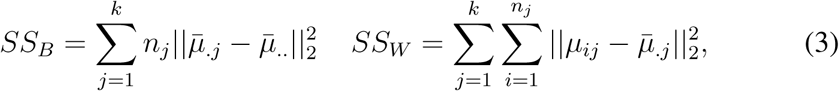

with:

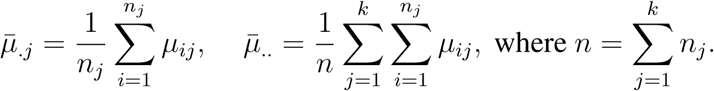

The statistic is known to approximately follow the *F* (*κ*(*k -* 1), *κ*(*n - k*)) distribution under the null hypothesis, where *κ* is a scaling constant obtained by two-cumulant approximation method (see (Zhang, 2013)). Due to low sample sizes for NGS experiments, the cumulant approximation is not reliable, thus in SRSFseq, we use the crude *F* (*k -* 1, *n - k*) distribution for the test statistic.

Equipped with this tool we move to application examples and performance evaluation of the new framework on RNA and DNA-seq data

### 2.2. Pre-processing of the raw data

To analyze the differences, first we have to perform the filtering step and obtain functional interpretation of sequencing over the exon locations (exon coordinates were obtained from the UCSC database (Kent et al., 2002), extracted in the form of a GTF file obtained from (Karolchik et al., 2004)). To do that we prvide R scripts (R Core Team, 2015), that assume as input the BAM files with mapped reads. Various software suites are available to transform the raw sequencing data into the designed format, we used samtools (Li et al., 2009) and bowtie2 (Langmead and Salzberg, 2012) for alignment against the human genome (HG19). In our analysis we are using the benchmark datasets described in (Trapnell et al., 2013). In each case the software parameters were set as described in the benchmark analysis of the same dataset.

As a result of this procedure we obtain a set of functions over shared reference region *G*, each function representing a different NGS experiment sample over different exon. Our goal is to compare the NGS experiments between conditions. A sample in the functional form from *j*-th condition is denoted 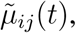, where *t* is the approximate position on in the common reference domain *G*. We assume that in each condition the observed filtered data 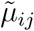 comes as a distortion of a unknown true density specific for the condition *j*, (denoted by *μ*_*j*_). We propose three ways of modelling intensities for detecting differences in NGS results, each model accounts for different types of distortions. In each model the intensities are normalized prior to performing analysis, so that 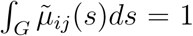. The models differ based on assumptions made on the source of variability between the intensities. In the discussion section we add one more model to show the possible extensions of the shape analysis framework.

### 2.3. SRSFseq: Base model (Base)

We propose a simple ANOVA - like setup for the observed, pre-processed density functions. The density representation of *i*-th sample in *j*-th condition is modelled as:

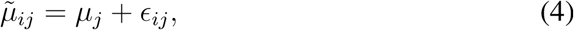

where *ϵ*_*ij*_ is a Gaussian stochastic process reflecting the noise in the data with 𝔼*ϵ*_*ij*_ = 0 and common covariance function *K*(*s, t*). *μ*_*j*_ is the base normalized density function of the *j*-th condition. *E*_*ij*_ is assumed to be pairwise independent. The density functions 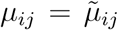 can be then directly applied in the test statistic described above.

The base model handles well obvious density differences as exemplified in Figure 3. As we show in section 3.1.1, even the simplest functional case proves to be useful in discovering new differential patterns.

**Figure 3:**
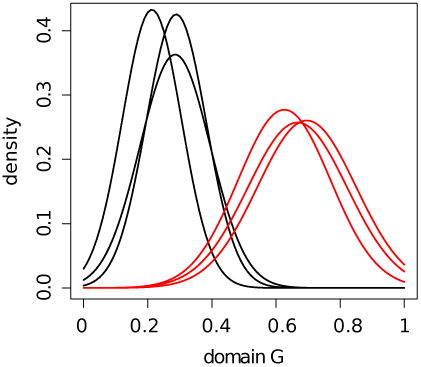
A simulated example of six samples of filtered density functions coming from *k* = 2 different condition with significantly different underlying true density functions *μ*_*red*_, *μ*_*black*_.

### 2.4. SRSFseq: noise removal (Shape and Energy preserving alignment)

Unfortunately, due to the low sample sizes of the NGS experiment, the filtering pre-processing step is very sensitive to the noise, when obtaining density functions. This may inflate the type I and type II errors in the test, due to misalignment in the filtered functions. To account for this issue we extend the analysis by an additional preprocessing step using the SRSF phase-amplitude separation method (Srivastava et al., 2011). We assume that the density functions might be distorted by domain shifts, which we will refer to as warping functions *γ*. Before conducting the analysis it is necessary to remove the warping noise, thus align the density functions. In our analysis we will look at two different type of distortions: shape preserving and energy preserving. Each model is capturing different aspects of the functional data and different ways to remove the noise.

#### 2.4.1. SRSFseq: Shape preserving noise removal (Shape)

We assume that observed warped density functions 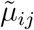 follow:

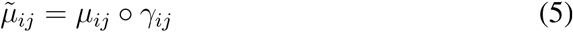

where *γ*_*ij*_ : *G* ↦*G* is an orientation preserving diffeomorphism corresponding to the warping noise. Using the phase-amplitude separation we are able to find optimal, shape preserving alignment 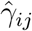 and use it to obtain undistorted intensities 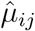

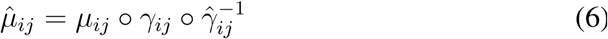

The warping *γ*_*ij*_ representing the phase noise, may have significant influence on the differential expression test, thus to properly evaluate the test statistic it is necessary to conduct the inference with the unwarpped density functions 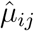. The process of aligning intensities by applying the composition with the warping function 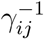 to 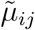 reduces the phase noise. The aligned intensities 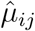 can be then used in the Functional ANOVA test statistic.

The aligned intensities are then modelled analogously to the base model

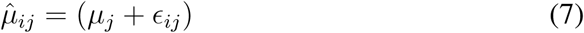

The advantage of the warping noise reduction in the model, is that we are able to eliminate misalignment between density functions, mistakenly inflating or deflating the values of the test statistic. To visualize the issue we have simulated six Gaussian curves which differ in amplitude or phase and compared them before and after alignment. In Figure 4 the curves were generated from two Gaussian intensities *μ*_*red*_ and *μ*_*black*_ significantly differing by the variance parameter. In panel A) the difference is not obvious from the point of view of the test as the *L*^2^ variability within each condition is comparable to the variability between each condition. After alignment the difference between conditions becomes apparent - panel B).

**Figure 4:**
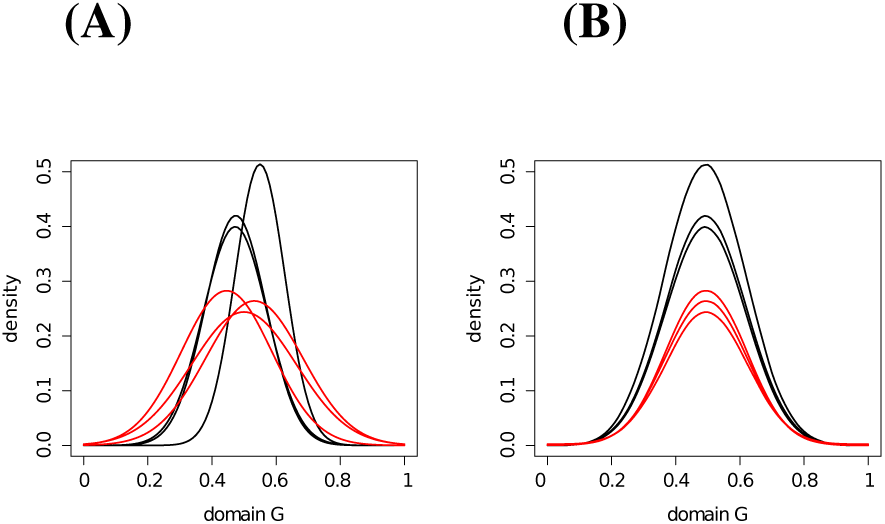
Six intensities generated from two conditions: red and black with significantly different true base density functions *μ*_*red*_, *μ*_*black*_. **A)** The unaligned raw intensities. **B)** The same density functions after the phase noise removal procedure.

The second scenario shown in Figure 5, corresponds to a problem where the random distortions accidentally drive the intensities to seem to be different. This can occur only by chance, but due to low samples sizes of the NGS experiment and the large number of genomic regions to be tested, such events have a non-negligible probability to occur, that has to be accounted for. As shown in Figure 5 six intensities were generated as distortions of the same Gaussian curve, but by accident three of them, consecutively generated, were shifted to the right (panel A). In such case the difference between those conditions could be falsely called significant. After alignment the groups of intensities seem indistinguishable (panel B).

**Figure 5:**
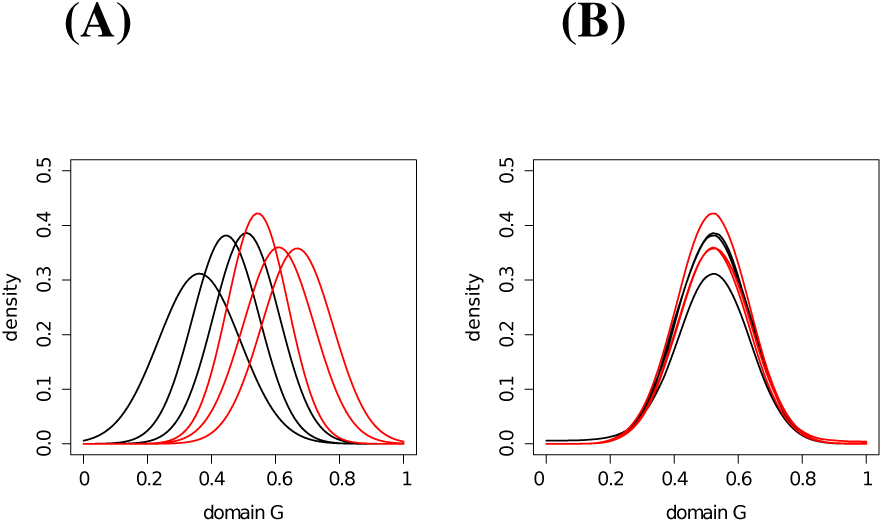
Six intensities generated from two conditions: red and black with same underlying true base density functions *μ*_*red*_ = *μ*_*black*_. **A)** The unaligned raw intensities. **B)** The same density functions after the shape noise removal procedure.

#### 2.4.2. SRSFseq: The energy preserving noise removal

In this model, as before we assume that the original density functions are distorted by a random warping *γ*_*ij*_. This time, however, the warping does not necessarily preserve the shape of the curve, but is constrained to maintain the energy (*L*^2^ norm). The intuition behind the energy-preserving model is the same as for the shape-preserving model, with the sole difference that the noise is introduced in Energy-preserving way. Accounting for the energy-preserving noise has the advantage over the shape-preserving noise, that it can cope with noise yielding significantly different shapes. The cost of it, however is that, the less constrained noise removal procedure (energy) may also accidentally remove critical information from the data.

We assume the following model for the distorted intensities:

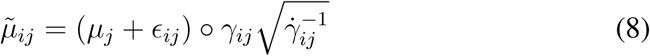

Using same SRSF phase-amplitude separation method (Srivastava et al., 2011) we obtain the optimal alignment between density functions 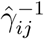 that preserve the energy norm of each curve.

The aligned model is then:

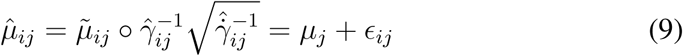

The obtained energy-aligned density functions 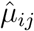 can be then further used in the functional ANOVA test statistic

## 3. Results

### 3.1. RNAseq expression analysis with Base, Shape and Energy models

To evaluate the information provided by SRSFseq we have compared the new functional models (denoted “Base”, “Shape” and “Energy”) with several other differential expression methods: Cufflinks, DESeq2, DEXSeq and Limma-voom (sections 2.3, 2.4 and 2.4.2) using published HOXA1 knock-out RNA-seq data (Trapnell et al., 2013). In our design a gene or exon is defined by the UCSC gene models. We treat each gene isoform as a separate gene and we call a gene differentially expressed if at least one of it’s exons shows a change in the density function.

The heat-maps in figure 6 show the dissimilarity between the SRSFseq and count-based methods. The heatmap entries indicate the number of genes called differentially expressed by both methods (row and column). For all of the new models, there is little overlap with count based methods. At the same time the SRSFseq methods have high overlaps. This is to be expected as SRSF normalization eliminates all count-based differences between samples. Interestingly, several genes identified specifically within our framework were related to developmental regulation, including COL1A1 and BAX, which have been previously implicated as targets of HOXA1 (Martinez-Ceballos et al., 2005; Zhang et al., 2003). What can be surprising at the beginning is that the SRSFseq methods report lower number of differentially expressed regions. This, however is also to be expected, as the differences in the shape of read density occur less frequently then count-based differences.

**Figure 6:**
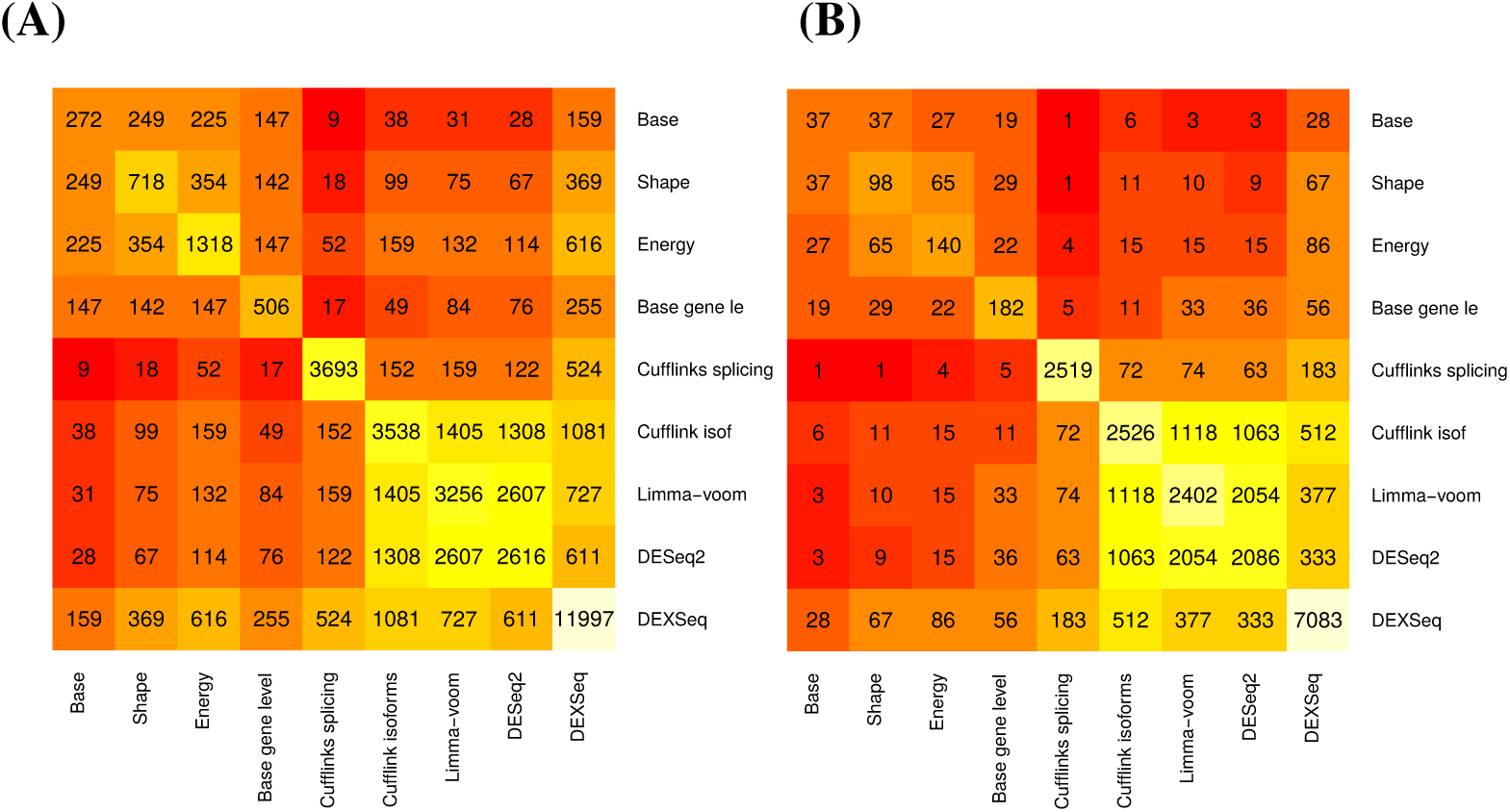
The heat-maps of the overlaps between lists of genes called differentially expressed by SRSFseq models (Base, Shape, Energy) and count based methods, using the significance level **(A)** *α*= 0.05, **(B)***α* = 0.01.

#### 3.1.1. New differential expression patterns uncovered

Figure (7AB) shows an example genes (uc001bvt.2, uc003vec.2), that displays a clear difference in filtered read densities, however this difference is not captured by any count-based methods because the overall counts at the genes do not significantly change (Cufflinks: p=0.229, DESeq2: p=0.908, Limma-voom: p=0.983). We have observed similar differential patterns in 272 genes called differentially expressed (*α*= 0.05) identified only by the base model. At the significance level of *α*= 0.01 30 out of 37 showed potential of exon overlap (Figure 7B), where the UCSC genome browser indicates an overlap between HBP1 and COG5 exactly on the region where the density functions differ.

**Figure 7:**
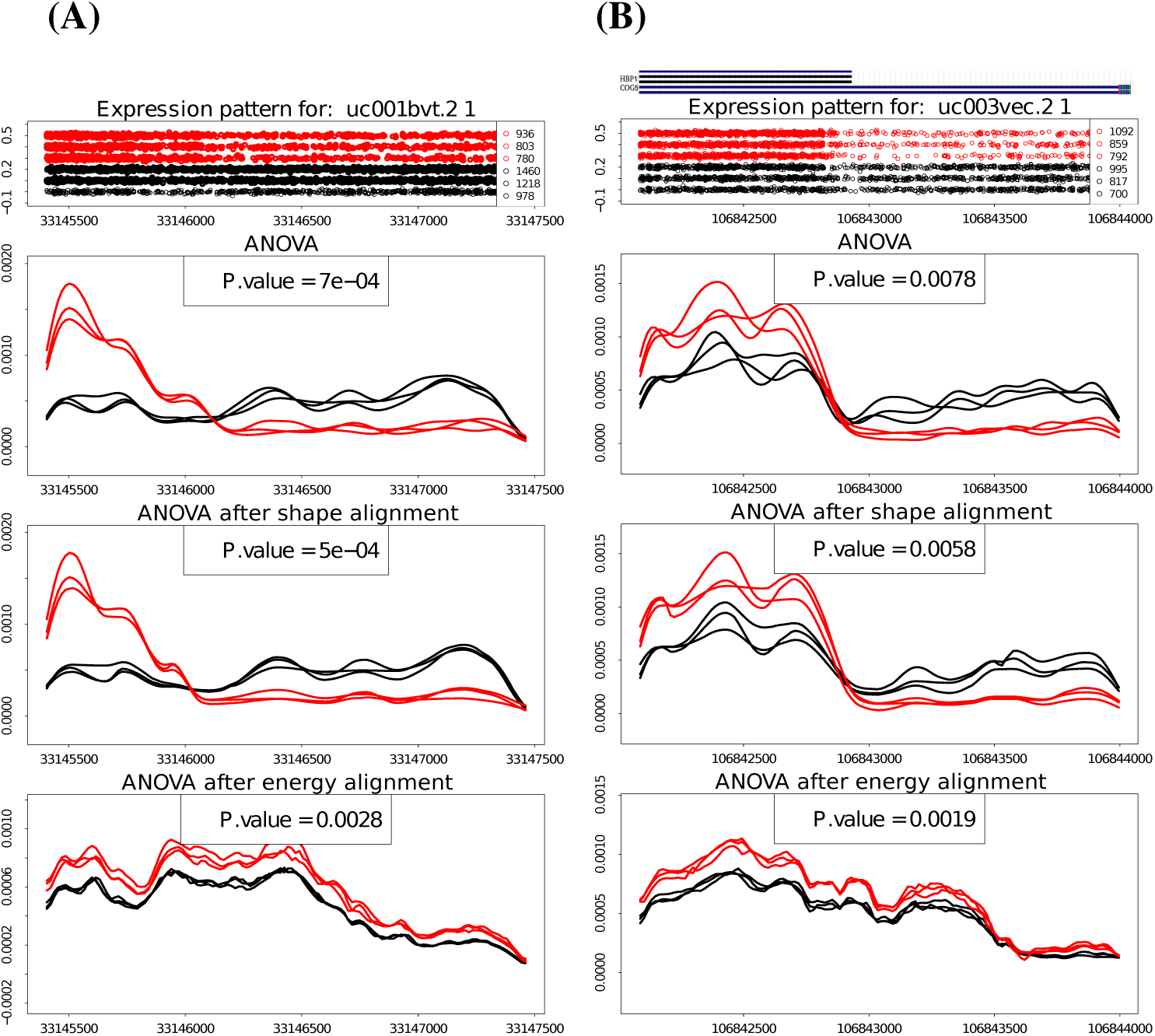
**(A)** Example of exonic region called differentially expressed by all three SRSFseq models, but not detected by any of the count-based methods, Two conditions: control (black), HOXA1 KO (red). Top panel: The point patterns over reference genome obtained by mapping first bp of each read, middle panel: filtered density functions, third panels: aligned density functions according to shape-preserving model, bottom panel: aligned density functions according to energy-preserving model. The p-values reported by other methods for the whole gene are: Cufflinks: 0.229, DESeq2: 0.908, Limma-voom: 0.983. **(B)** Similar example, but for convenience we provide the UCSC genome browser screen-shot for the region on top of the main figure. The comparison with genome browser indicates that the new differential patterns detected by SRSFseq can be explained by the current knowledge about gene location. The p-values reported by other methods: Cufflinks: 0.077, DESeq2: not reported, Limma-voom: not reported.

#### 3.1.2. Advantage of the shape- and energy- preserving noise removal

Accounting for the phase variability is designed to improve the results of the new method by controlling for variability in read distributions that would result in false positives and affect the statistical significance of the truly differentially expressed genes that were not detected by the base model alone (nor by any count based methods) (Figure 8).

**Figure 8:**
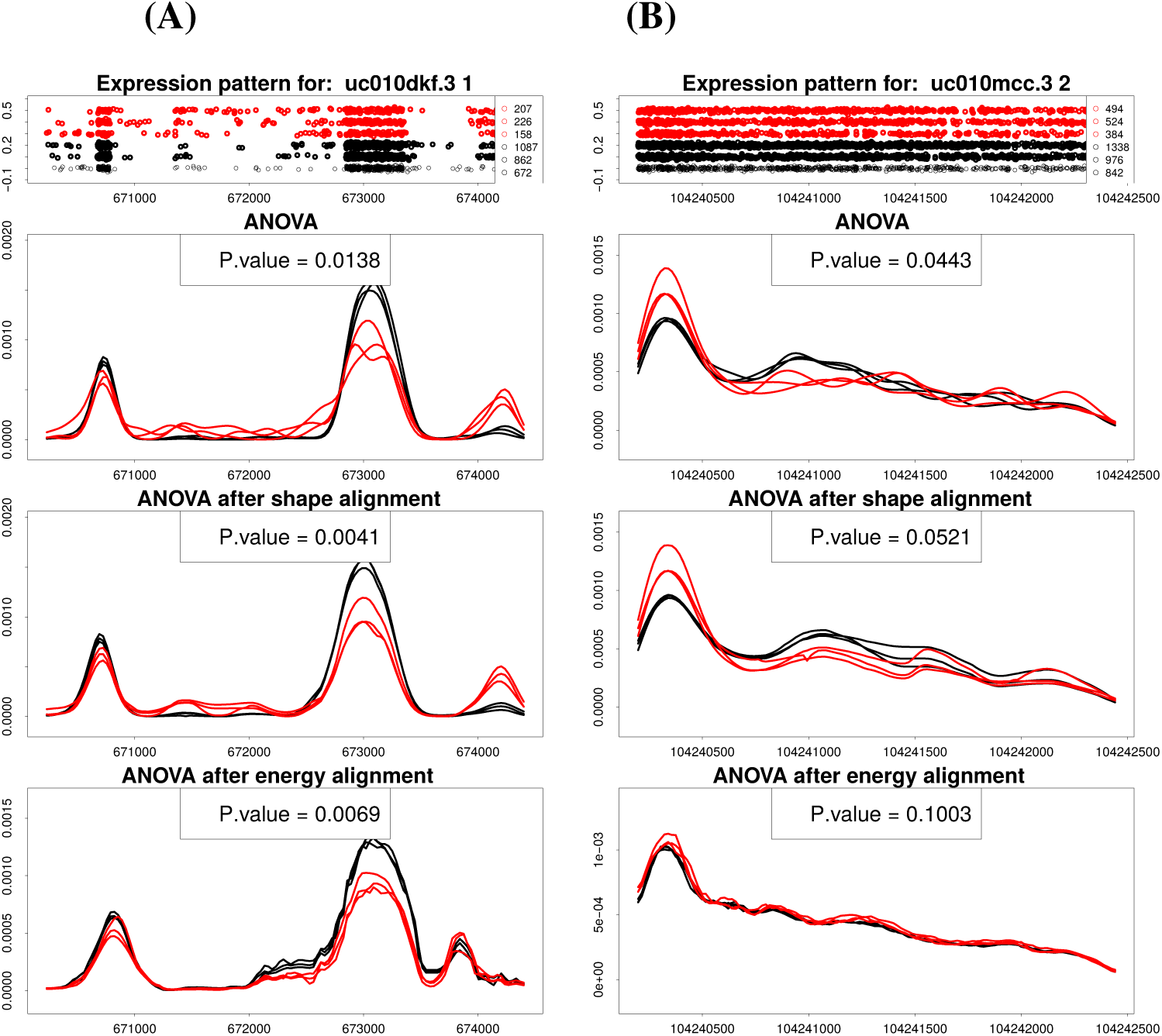
**(A)** Example of an exonic region called differentially expressed on the significance level of *α*= 0.01 only after the shape noise is removed. Two conditions: control (black), HOXA1 KO (red). Top panel: The point patterns over the reference genome obtained by mapping first base pair of each read. Middle panel: filtered density functions. Bottom panel: aligned density functions. **(B)** Example of an exon region that was called differentially expressed by the base model, but lost significance after applying the shape-noise removal procedure. The sum of square distances were inflated due to the noise. Two conditions: control (black), HOXA1 KO (red). Top panel: The point patterns over the reference genome obtained by mapping the first base pair of each read. Middle panel: The observed filtered density functions. Bottom panel: The aligned density functions.

Figure 8A shows an example gene, X, where noise removal procedure can improve the differential expression detection. The alignment of the density functions helps reduce the *SS*_*W*_ component of the test statistic, which previously was keeping the statistic result below the significance level *α*= 0.01. Figure 8B shows an example gene, X, where the noise removal procedure increases the p-value above the significance level of *α*= 0.05, consistent with a lack of strong evidence for the differential expression based on read densities alone.

In both instances (A,B) the noise removal improved the results by either capturing a False Positive, False Negative

Which noise removal is superior? In the Figure 9 we highlight that, although the noise removal is desirable and overall performance improves when compared with base model, neither of the proposed alignments proves to be significantly better than the other. Figures 9A,B show the advantage of the energy-preserving alignment over the shape-preserving alignment model by reducing the type I and II errors. The third panel however (Figure 9C) emphasizes that, even though energy-preserving alignment seems to perform better, it’s relatively weak constraints (constant energy or *L*^2^ norm), may cause information loss after noise removal. In particular the energy-preserving aligning method won’t distinguish between density functions that have same energy even if their shapes are significantly different, as seen on the bottom panel of the figure.

**Figure 9:**
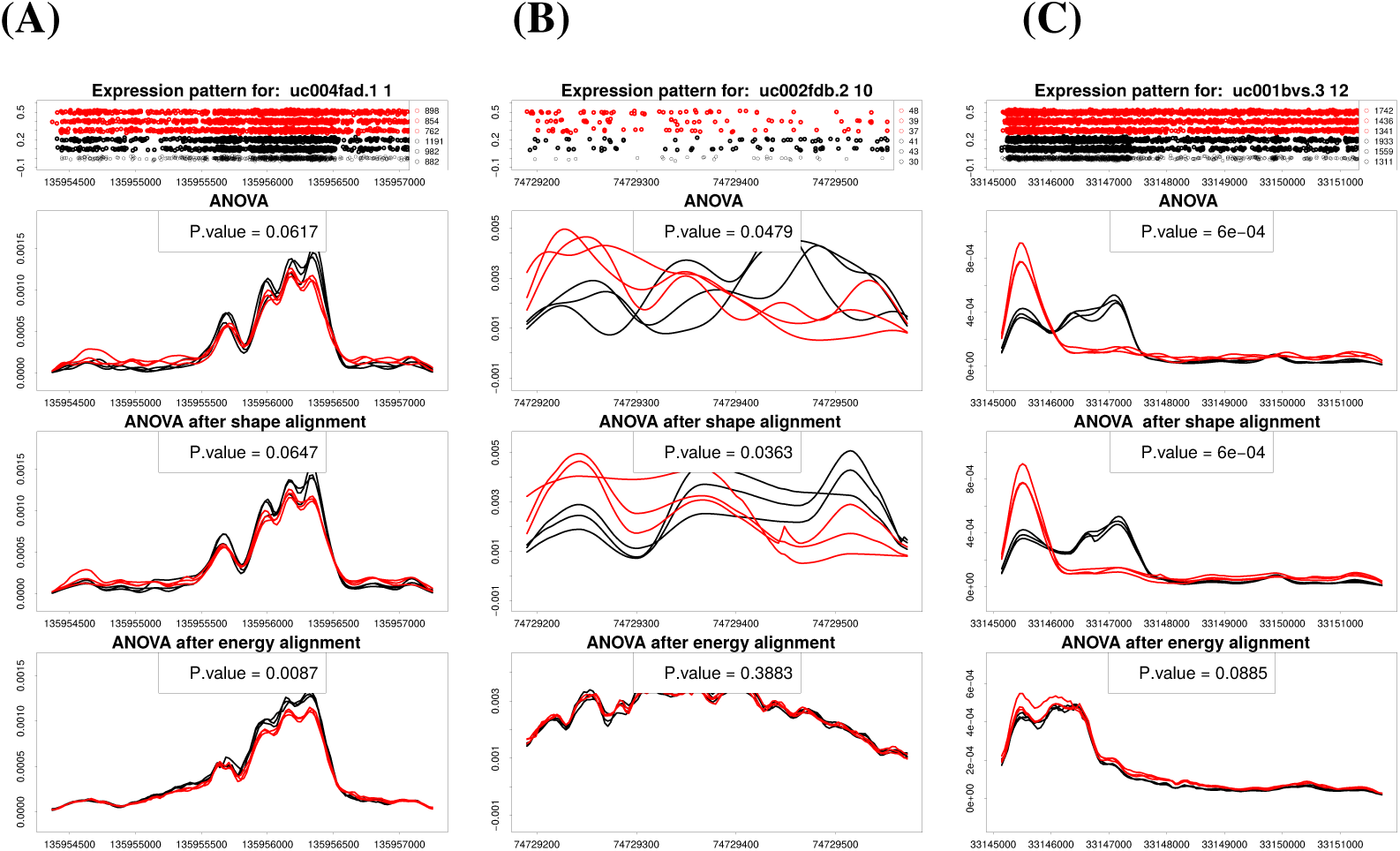
**(A)** Energy-preserving noise removal improves detection of differences comparing to shape-preserving alignment and base model, by capturing a false positive. **(B)** Energy-preserving noise removal improves detection of differences comparing to shape-preserving alignment and base model, by capturing a false negative. **(C)** Energy-preserving noise removal causes loss of information and fails to detect a significant difference between expression patterns. This difference is successfully captured by the shape-preserving alignment and base model.

### 3.2. Misalignment as differences in activity patterns

The proposed functional framework describes a novel way of modelling the genomic distribution of reads by identifying the differences in the shape of the read density functions between experimental conditions. To show the potential of SRSFseq we exemplify how our generative models (energy -and shape-preserving alignment), can be extended, by alternating roles of the aligning components in the model (*γ* functions) In the new setting we assume that the observed intensities 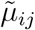 arrive as condition – specific shape changes *γ*_*j*_ of the same true base density μ and test whether the shape changes *γ*_*j*_ are significant between conditions The aligning functions are *γ* no longer recognized as noise – they now carry potentially significant information

In short, the we assume that:

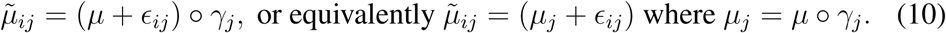

and we aim to test the null hypothesis of no difference between the shape changes between any two conditions *j*_1_, *j*_2_: *H*_0_ : *γ*_1_ = *γ*_2_ = *…* = *γ*_*k*_. Or alternatively: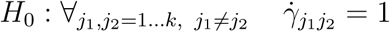, where 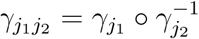.

As *γ*_*j1j2*_ can be estimated by the phase-amplitude separation algorithm (Srivastava et al., 2011), we can measure the magnitude of local shifts between conditions, the location of those shifts, or quantify the differences between aligned patterns. As a consequence of the design, this particular model has little application in direct exon-level expression analysis, because there is not a known biological interpretation for RNA-seq read densities to be consistently shifted. It opens, however, the possibility to analyze shifts or shape changes in genomic data problems, for which very few approaches are currently available, e.g. differences in the positions of nucleosomes (Chen et al., 2013; He et al., 2010; Meyer et al., 2011; Fu et al., 2012).

Below we present a work-in-progress example application of the alternative model design to detect differences in nucleosome positions between experimental conditions (”shifts”) near the transcription starting sites (TSS). The motivation for such analysis is to test the hypothesis that the nucleosome shifts near TSS is associated with the regulation of gene expression. Few approaches have been developed to identify differences in nucleosome positions that are applicable to large eukaryotic genomes such as human (cite danpos2 and others based on emailed list). Using MNase-seq data, we are able to perform the phase-amplitude analysis and estimate the optimal warping between the density curves in two different cell types (lymphocytes and fibroblasts). The Figure 10A shows an example piece of information that can be easily decoded by the model. The figure presents the density functions with their aligned versions combined with the location and amount of the shift (in bp per TSS). We can observe a clear shifting of the density peaks towards the right end of the graphs, which have a confirmation in high absolute values of the 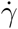 graphed in the top panel.

**Figure 10:**
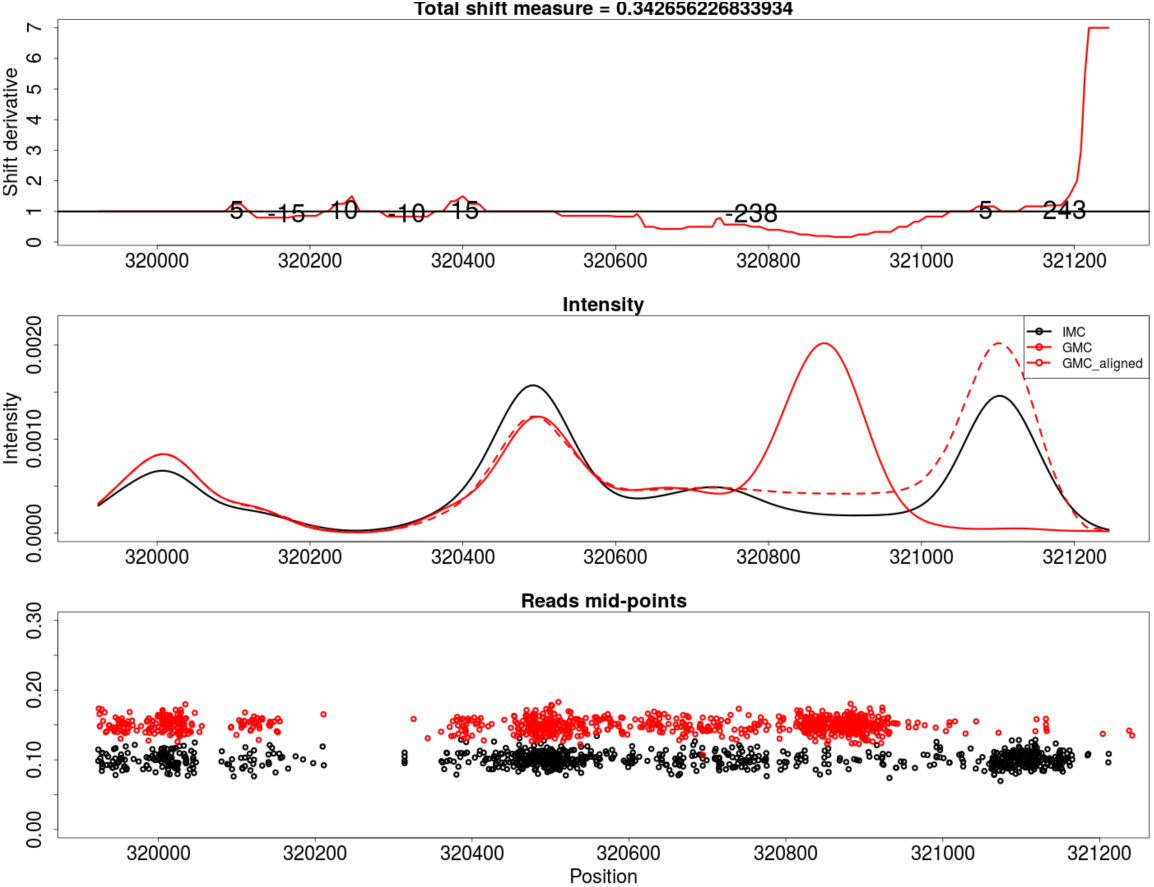
**(A)** Shift detection near the TSS regions. Comparing the DNASeq filtered density between two conditions: control (Black) and nucleosome shift (Red). Bottom panel shows the endpoints positions of the mapped reads on the reference genome, Middle panel shows the filtered density functions, with the additional dashed red line reflecting the optimal alignment of the red density to the black density. Top panel shows the relative changes in density domain with respect to the control density. Red curve above the black horizontal line reflects the shifts toward 5’ end; below, towards 3’ end. Area between black and red curves reflect the amount of shifting needed.

This model will be further developed for application to MNase-seq data and other chromatin structure data where shifts in read densities can be used to infer differences in chromatin structure.

## 4. Discussion

We have proposed a new framework (SRSFseq) of investigating the NGS data through the functional interpretation. We have shown that the new approach can be successively used in analysing NGS outcomes and uncover information not possible to decode with the state-of-the-art methods. We have equipped the SRSFseq with two functional noise removal procedures improving the type I and type II errors. Interestingly, if we performed same analysis on whole spliced genes, instead of on separate exons, we still obtain significantly different gene lists called differentially expressed comparing to both exon-level SRSFseq and all count-based methods. The results can be seen on Figure 10B.

We have shown the flexibility of SRSFseq and have given the examples of how it can be tuned to address the experimental questions. We have proposed an alternative application of the framework that aims do detect changes the shape of read density and exemplified it on nucleosome shift detection problem.

In the filtering step in this article we have used the simple kernel density estimator, however we would like to point out, that other density estimation techniques can be used depending on particular applications, especially when one needs to account for over-dispersion of read counts or read clustering problem.

We would like to emphasize that the new framework in case of the RNAseq data, due to it’s normalization procedure, won’t detect gene-wide count based differences. As such, it should not be viewed as a replacement for methods that detect global differences in gene expression, but can be effectively used to supplement their results where overlapping genes result in false negatives.

## 5. Software

Software in the form of R script, used to obtain results in this article is available in a reproducible and adjustable way, on public Github repository: https://github.com/FSUgenomics/SRSFseq

## References

Anders, S., Reyes, A., Huber, W., 2012. Detecting differential usage of exons from rna-seq data. Genome research 22 (10), 2008–2017.

Chen, K., Xi, Y., Pan, X., Li, Z., Kaestner, K., Tyler, J., Dent, S., He, X., Li, W., 2013. Danpos: dynamic analysis of nucleosome position and occupancy by sequencing. Genome research 23 (2), 341–351.

Fu, K., Tang, Q., Feng, J., Liu, X. S., Zhang, Y., 2012. Dinup: a systematic approach to identify regions of differential nucleosome positioning. Bioinformatics 28 (15), 1965–1971.

Hardcastle, T. J., Kelly, K. A., 2010. bayseq: empirical bayesian methods for identifying differential expression in sequence count data. BMC bioinformatics 11 (1), 422.

Hayer, K. E., Pizarro, A., Lahens, N. F., Hogenesch, J. B., Grant, G. R., 2015. Benchmark analysis of algorithms for determining and quantifying full-length mrna splice forms from rna-seq data. Bioinformatics, btv488.

He, H. H., Meyer, C. A., Shin, H., Bailey, S. T., Wei, G., Wang, Q., Zhang, Y., Xu, K., Ni, M., Lupien, M., et al., 2010. Nucleosome dynamics define transcriptional enhancers. Nature genetics 42 (4), 343–347.

Karolchik, D., Hinrichs, A. S., Furey, T. S., Roskin, K. M., Sugnet, C. W., Haussler, D., Kent, W. J., 2004. The ucsc table browser data retrieval tool. Nucleic acids research 32 (Suppl 1), D493–D496.

Kent, W. J., Sugnet, C. W., Furey, T. S., Roskin, K. M., Pringle, T. H., Zahler, A. M., Haussler, D., 2002. The human genome browser at ucsc. Genome research 12 (6), 996–1006.

Langmead, B., Salzberg, S. L., 2012. Fast gapped-read alignment with bowtie 2. Nature methods 9 (4), 357–359.

Law, C. W., Chen, Y., Shi, W., Smyth, G. K., 2014. Voom: precision weights unlock linear model analysis tools for rna-seq read counts. Genome Biol 15 (2), R29.

Leng, N., Dawson, J. A., Thomson, J. A., Ruotti, V., Rissman, A. I., Smits, B. M., Haag, J. D., Gould, M. N., Stewart, R. M., Kendziorski, C., 2013. Ebseq: an empirical bayes hierarchical model for inference in rna-seq experiments. Bioinformatics, btt087.

Li, H., Handsaker, B., Wysoker, A., Fennell, T., Ruan, J., Homer, N., Marth, G., Abecasis, G., Durbin, R., et al., 2009. The sequence alignment/map format and samtools. Bioinformatics 25 (16), 2078–2079.

Li, Y., Rao, X., Mattox, W. W., Amos, C. I., Liu, B., 2015. Rna-seq analysis of differential splice junction usage and intron retentions by dexseq. PLoS one 10 (9), e0136653.

Love, M. I., Huber, W., Anders, S., 2014. Moderated estimation of fold change and dispersion for rna-seq data with deseq2. Genome biology 15 (12), 1–21.

Martinez-Ceballos, E., Chambon, P., Gudas, L. J., 2005. Differences in gene expression between wild type and hoxa1 knockout embryonic stem cells after retinoic acid treatment or leukemia inhibitory factor (lif) removal. Journal of Biological Chemistry 280 (16), 16484–16498.

Meyer, C. A., He, H. H., Brown, M., Liu, X. S., 2011. Binoch: binding inference from nucleosome occupancy changes. Bioinformatics 27 (13), 1867–1868.

R Core Team, 2015. R: A Language and Environment for Statistical Computing. R Foundation for Statistical Computing, Vienna, Austria. URL https://www.R-project.org/

Srivastava, A., Wu, W., Kurtek, S., Klassen, E., Marron, J., 2011. Statistical analysis and modeling of elastic functions. arXiv preprint arXiv:1103.3817.

Trapnell, C., Hendrickson, D. G., Sauvageau, M., Goff, L., Rinn, J. L., Pachter, L., 2013. Differential analysis of gene regulation at transcript resolution with rna-seq. Nature biotechnology 31 (1), 46–53.

Trapnell, C., Roberts, A., Goff, L., Pertea, G., Kim, D., Kelley, D. R., Pimentel, H., Salzberg, S. L., Rinn, J. L., Pachter, L., 2012. Differential gene and transcript expression analysis of rna-seq experiments with tophat and cufflinks. Nature protocols 7 (3), 562–578.

Zhang, J.-T., 2013. Analysis of variance for functional data. CRC Press.

Zhang, X., Zhu, T., Chen, Y., Mertani, H. C., Lee, K.-O., Lobie, P. E., 2003. Human growth hormone-regulated hoxa1 is a human mammary epithelial oncogene. Journal of Biological Chemistry 278 (9), 7580–7590.

